# Anti-inflammatory assessment of zinc oxide nanoparticles mediated *Aframomum citratum* (C. Pereira) K. Schum (Zingiberaceae) in Wistar rats

**DOI:** 10.1101/2024.12.15.628600

**Authors:** Francois Eya’ane Meva, Denise Murielle Nga Essama, Edvige Laure Nguemfo, Hans-Denis Bamal, Agnes Antoinette Ntoumba, Phillipe Belle Ebanda Kedi, Thi Hai Yen Beglau, Alex Kevin Tako Djimefo, Annie Guilaine Djuidje, Geordamie Chimi Tchatchouang, Chick Christian Nanga, Gildas Fonye Nyuyfoni, Armel Florian Tchangou Njimou, Danielle Ines Madeleine Evouna, Armel Ulrich Mintang Fongang, Christoph Janiak

**Author notes:** Corresponding author: Francois Eya’aneMeva.

## Abstract

**Introduction:** Zinc oxide nanoparticles (ZnONPs) have been synthesized using a wide range of techniques, including green chemistry, because of their versatility, cost effectiveness, and environmentally friendly nature, offering thereby interesting and inexpensive therapeutic options. This study aimed to develop zinc oxide nanoparticles as an anti-inflammatory agent using *Aframomum citratum* seed extract.

**Methodology:** ZnONPs were prepared by the reaction between zinc nitrate and an alkalineaqueous extract of *A. citratum* seeds. The isolated nanoparticles were then characterized using UV-Vis, FTIR, SEM/EDX, PXRD and TEM techniques. The toxicological profile was assessed at a limited dose of 2000 mg/kg in rats, and methods for heat denaturation of egg albumin, stabilization of red blood cell membranes and inhibition of carrageenan-induced plantar oedema were studied to assess anti-inflammatory properties.

**Results:** The formation of ZnONPs was observed by a color change and the appearance of the plasmon resonance peak at 360 nm in the UV-Vis spectrum while FTIR confirmed the presence of secondary metabolites; SEM confirmed the presence of multiform aggregates, and TEM visualize point like particles. EDS confirmed the presence of Zn atoms within the synthetized material. The toxicological profile studied showed no harmful signs; zinc oxide nanoparticles synthesized from *A. citratum* seed extract showed high inhibition percentages of 86 (1mg/mL); 77 (0.6mg/mL) and 79(1mg/mL) when subjected to inhibition of heat-induced egg albumin denaturation, red cell membrane stabilization and oedema induction by carrageenan respectively, not significatively different compared with diclofenac sodium as positive controls.

**Conclusion:** Zinc oxide nanoparticles synthesized and characterized from *A. citratum* seed extract act as a potent anti-inflammatory agent and are devoid of acute oral toxicity.

## Introduction

According to the World Health Organization (WHO), three out of five people die every year from inflammation-related pathologies. Inflammation occurs when the integrity of an organism’s morphological or biological constants is threatened. It occurs during infections or aggression by physical and chemical agents. Inflammation is manifested by symptoms such as oedema, pain, heat, or fever [1, 2]. Current treatments for inflammatory diseases based on non-steroidal anti-inflammatory drugs (NSAIDs) and steroids may have limitations on some clinical events, such as complications of the gastrointestinal tract, a history of heart disease, ulcers, and bleeding. In addition, their cost and poor accessibility are major obstacles in a context of countries with limited resources.

For centuries, medicinal plants have had an established therapeutic value for a variety of human ailments. Their effects derive from the secondary metabolites of their extracts [2, 3]. The application of nanotechnology to herbal medicines could lead to the development of nanoproducts with enhanced solubility, bioavailability, and pharmacological activity [3]. Zinc is an essential element for human life. Studies on zinc oxide show that it is biodegradable and pharmacologically interesting. The powder form is incorporated in baby powders and barrier creams to treat or prevent a variety of skin conditions including dermatitis. It can be mixed with iron oxide to form calamite lotion used as topical skin treatment, or castor oil to form emollient and astringent. Also, due to its antibacterial properties zinc oxide is incorporated in various oral care products such as toothpaste, mouthwash to prevent tartar and plaque formation [4–5].Its use could lead to exiting synergistic situation when combined with plant extracts.

*A. citratum* is a species found in the Guinean forests of Africa. It is also distributed in Nigeria, Cameroon, and Gabon[6]. *A. citratum* is considered a hard-shelled fruit and a non-wood forest product, used as a spice and in many traditional pharmacopoeia recipes. It is heat-treated before being ground in the recipe for “*Mbongo Tchobi*”, a spicy black sauce native to the central and coastal regions of Cameroon. Oral administration of hydroethanolic extracts of *A. citratum, A. danielli* and *A. aulacocorpus* reduced the exponential weight gain induced by the atherogenic diet and was accompanied by a significant (p<0.05) reduction in high-density lipoprotein (HDL) cholesterol.

With the aim of proposing options for the management of inflammatory pathologies while protecting biodiversity, this study presents the synthesis of zinc oxide nanoparticles (ZnONPs) using *A. citratum* seed extract, their acute toxicological profile, and assessment of their anti-inflammatory activity *in vitro* and *in vivo*.

## Experimental

### Collection, authentication, and preparation of extract

*A. citratum* fruits were harvested in Ngoulemakong (N°3 04ʹ 60.00ʺ; E°11 25ʹ 59.99ʺ) 50 km from Ebolowa in the South Region of Cameroon and authenticated at the Cameroon National Herbarium as identical to specimen number 37736/HNC. After harvesting, the fruits were washed with tap water and then with distilled water to remove all contaminants. The seeds were removed from the pulp, dried for two weeks, away from light, and then pulverized to obtain the powder (Figure 1). An infusion was then made at 60°C for 10 minutes by dissolving 75g of powder in 750mL of distilled water (mass/volume ratio=1/10 in g/mL). The mixture was filtered through Nᵒ1 Whatman filter paper and concentrated by removing residual solvent in an oven at 40 °C. The extraction yield was then determinedas a percentage of the dry extract in the infused powder.

**Figure 1.**
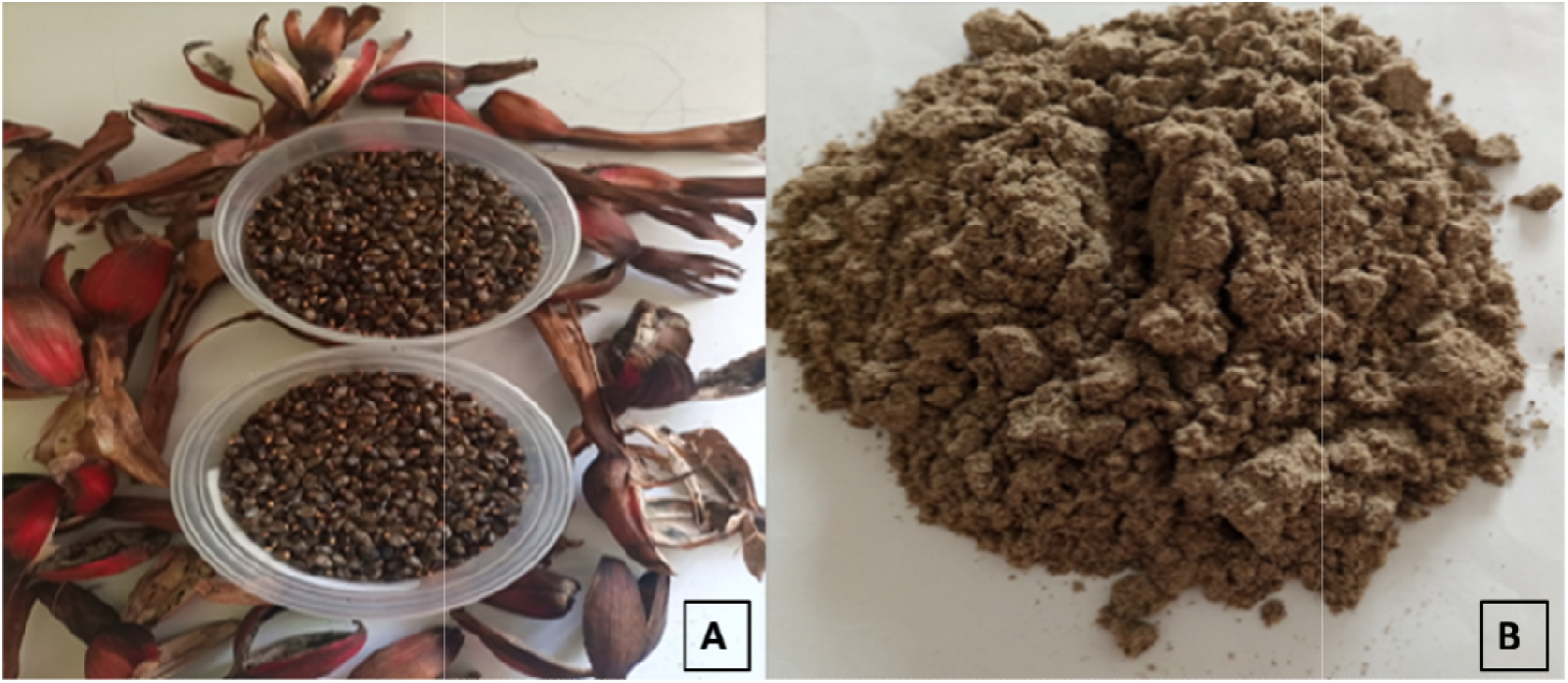
Dried fruits and seeds (A), and powder (B) from the seeds of *Afromomum citratum*

### Phytochemical study

Several tests were carried out to determine the presence or absence of various families of secondary metabolites in the plant extract. These included polyterpenes and sterols, using the Liebermann Burchard reaction, cardiotonic glycosides according to Shivakumar and collaborators, as described elsewhere [7], alkaloids by the Dragendorff, Valser-Mayer reaction [8], tannins according to Nemlin and Brunel, 1995 [19]. Polyphenols, flavonoids, and coumarins were determined using methods previously reported in the literature [10, 11].

### Synthesis of zinc oxide nanoparticles

A solution of 0.2 mol/Lzinc nitrate hexahydrate and a solution of 2 mol/L NaOH were prepared. 30 mL of *A. citratum* aqueous extract was mixed with 70 mL zinc nitrate hexahydrate, and the solution was stirred for 10 min. The pH of the reaction mixture was adjusted by the dropwise addition of 2 mol/L NaOH at pH 12. Synthesis was monitored by a visual observation of color change of the colloid suspension.UV-Vis spectrophotometric scanning between 300 and 600nm were accessed to find the plasmon resonance, indicator of the formation of ZnONPs. These nanoparticles were centrifuged at 4000 rpm for 20 min and washed twice with distilled water and then ethanol to remove impurities. The final product in the form of a white powder material was dried overnight at 50°C [12].

### Ultraviolet-visible characterization (UV-Vis)

Ultraviolet-Visible spectroscopy was monitored on a P9 double beam spectrophotometer from VWR. Measurements were made between 200 and 800 nm.

### Fourier-transformed infrared spectroscopy (FTIR)

Infrared spectroscopy was conducted using a Bruker Tensor 37 with attenuated total reflection, ATR unit by scanning between 600-4000 cm^−1^.

### Powder X-ray diffraction (PXRD)

The PXRD diffraction of the nanoparticles was performed using a Bruker D2 Phaser powder diffractometer (Cu K-Alpha1 [Å] 1.54060, K-Alpha2 [Å] 1.54443, K-Beta [Å] 1.39225) by preparing a thin film on a low-background silicon sample holder; 30 min measurement time, 2 theta range 5°– 80°.

### Scanning electron microscopy (SEM) and energy-dispersive X-ray spectroscopy (EDX) investigations

Scanning electron microscopy (SEM) was carried out with a Jeol JSM−6510LV QSEM Advanced electron microscope with a LaB_6_ cathode at 20 kV equipped with a BrukerXflash 410 (Bruker AXS, Karlsruhe, Germany) silicon drift detector for energy-dispersive X-ray spectrometric (EDX) analysis. The samples containing ZnONPs were sputtered with gold by using a JEOL JFC-1200 Fine Coater.

### Transmission electron microscopy (TEM)

Electron micrographs of the nanoparticles were recorded using a JEOL JEM-2100 Transmission Electron Microscope at 200 kV with a TVIPS F416 camera system.

### Animal and ethical considerations

Healthy adult female Wistar albino rats weighing between 120 – 180 g were obtained from the Animal Laboratory of the Faculty of Medicine and Pharmaceutical Sciences, University of Douala, Cameroon. The animals were sorted randomly in standard polypropylene cages in groups of three and kept under standard conditions of temperature (24 ± 2°C) and light (approximately 12 h/12 h light/dark cycle) with free access to standard laboratory diet and tap water *ad libitum*. An ethical clearance Nr 3544 CEI-Udo/03/2023T was obtained from the Institutional Ethics Committee for Human Health Research at the University of Douala, Cameroon.

### Assessment of acute oral toxicity

The acute toxicity test was carried out following OECD guidelines N°425 pertaining to acute toxicity [13] at limit dose of 2000 mg/kg. Briefly, two batches of 3 female rats were randomized. Batch 1 received the synthesized nanoparticles (ZnONPs) at 2000 mg/kg, while Batch 2 served as control and received 10 mL/kg distilled water. The rats were weighed and fasted with free access to water 24 hours before the start of the test. Rats were observed individually for 14 days, with particular attention paid to the first 30 minutes, then at 4h, 8h, and after 24 h post-administration. Clinical observations were tremor, convulsion, salivation, diarrhea, lethargy, sleep, coma, skin, hair, eyes, respiratory tract, and changes in skin, hair, and eye mucosa. Rats were weighed every two days, and their weights were recorded. On day14, the rats were sacrificed by inhalation formalin anesthesia in a closed jar. Blood was collected after incision of the carotid artery and later analyzed by creatinine urea test and alanine and aspartate aminotransferases (ALT/AST) test for the determination of biochemical parameters providing information on renal and hepatic functions, respectively. Some organs such as kidneys, liver, spleen, lungs, and heart were removed after the dissection for analysis of biochemical parameters, then weighed and their relative masses were calculated.

### Heat-induced egg albumin denaturation assay

The anti-inflammatory activity of ZnONPs with *A. citratum* extract was assessed *in vitro* using the heat-induced egg albumin denaturation assay, as reported elsewhere [14]. Briefly, the reaction media (5 mL) consisted of 2.8 mL of phosphate buffer saline (PBS) adjusted to pH = 6.4 with HCl and 0.2 mL of egg albumin, to which 2 mL of tested substances were added. The aqueous extract and the ZnONPs were tested at different concentrations (200, 400, 600, 800 and 1000 µg/mL each). Diclofenac at the same concentrations served as reference while distilled water served as control. All the mixtures were incubated at 37°C for 15 min, then gradually heated up to 70°C and kept at that temperature for 5 min before allowing them to cool down for 20 min. The absorbance was measured for each mixture with a spectrophotometer at 660 nm and the reaction percentage was calculated using the following formula 1:

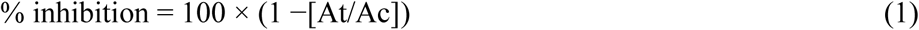

with;

At: Absorbance of sample (test)
Ac: Absorbance of the control

### Preparation of erythrocyte suspension

Blood samples were collected from Wistar rats in heparinized tubes and centrifuged at 3000 rpm for 10 min. The supernatant was then removed, and the red blood cell pellet was washed three times with a PBS solution (pH= 7,2 - 7,4) and centrifuged each time at the same speed for 5 min. The volume of red blood cells was measured, and a 5% suspension was prepared by diluting into PBS.

### Erythrocyte membrane stabilizationassay

The test is based on the effect of ZnONPs and aqueous extract of *A. citratum*on erythrocyte membrane stabilization, after induction of hemolysis by a hypotonic solution combined with elevated temperature following a method described elsewhere, with slight modification. In different test tubes, 0.5mL of ZnONPs, aqueous extract,or diclofenac sodium were respectively dissolved in 0.9% NaCl, then 1.5mL of phosphate buffer (pH=7.4) and 2mL of hyposaline solution (0.36% NaCl) were mixed and incubated at 37°C for 20 min. A volume of 0.5mL of erythrocyte suspension (5%) was then added to each tube, followed by incubation at 56°C for 30 min. The tubes were centrifuged at 3000 rpm for 5 min, and then the absorbance of the supernatant was read at 560nm. The control consisted of a mixture of 2mL hyposaline solution, 2mL PBS, 0.5mL erythrocyte suspension and 0.5mL distilled water. The hemolysis inhibition rate was calculated according to the following equation (1)

### Evaluation of anti-inflammatory activity *in vivo*

The anti-inflammatory effect of ZnONPs with *A. citratum* extract was assessed *in vivo* on carrageenan-induced rat paw edema models, as described by other authors, with some modifications [15]. Briefly, five groups (n = 5) of Wistar rats were formed and the animal werefasted 12h prior to the experimentation. An initial volume V_0_ of the left hind paw of each rat was measured using a plethysmometer, before administering the different solutions orally. Three groups received the ZnONPs with*A. Citratum* extract at 600, 800 and 1000 μg/kg, respectively. One group received diclofenac (10 mg/kg) and served as positive control. A final group received distilled water (10 mL/kg) and served as negative control. A single sub-plantar injection of 100 μL of 1% carrageenan (1% carrageenan suspended in 0.9% NaCl) was performed into the left hind paw of each animal, 30 min after administration of the solutions. The volume of the left hind paw of each rat was then measured at 30 min and every hour for 6 hours. The percentage of inhibition at each time of measurement was obtained using the following formula (2):

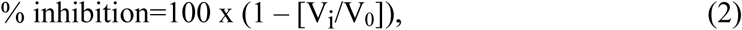

Where V_0_ stands for edema volume in control and V_i_ the edema volume in the test groups at different hours

### Statistical analysis

Quantitative data was analyzed using GraphPad Prism for Windows software (version 9.1) and presented as the mean ± standard error of the mean. The results were compared by two-way ANOVA followed by Tukey’s multiple comparison at the 95% threshold.

## Results

### Plant extract studies

The pulverized seeds from *A. citratum* (75 g) were used for extraction by infusion into 750 mL of distilled water at 60°C, and the mixture was filtered and dried. The extract obtained was a 7 g brown colored paste, indicating a 9 % extraction yield. The qualitative phytochemical screening of *A. citratum*’s aqueous extract revealed the presence of some of the secondary metabolites tested for, namely phenolic compounds (flavonoids, tannins, anthraquinones) and terpenoids (saponins). Nevertheless, some classes were absent, such as coumarins, anthocyanins, triterpenes, sterols and reducing sugars as primary metabolites (Table 1).

**Table 1.** Phytochemical content of the aqueous extract of *Afromomumcitratum* seed.

| Tested metabolites | Results |
| --- | --- |
| Alkaloids | + |
| Phenolics | + |
| Flavonoids | + |
| Tannins | + |
| Coumarins | + |
| Anthraquinones | + |
| Anthocyanins | - |
| Sterols and terpenes | + |
| Triterpenes | - |
| Saponins | - |
| Reducing sugars | - |

(+) present; (-) absent

### Synthesis of zinc oxide nanoparticles

#### Visual observation

Figure 2 below shows the change in color of the brown-colored aqueous extract of *A. citratum* after the addition of zinc nitrate hexahydrate to give a pale white dispersion; the opaque whitening marks the formation of zinc oxide nanoparticles.

**Figure 2.**
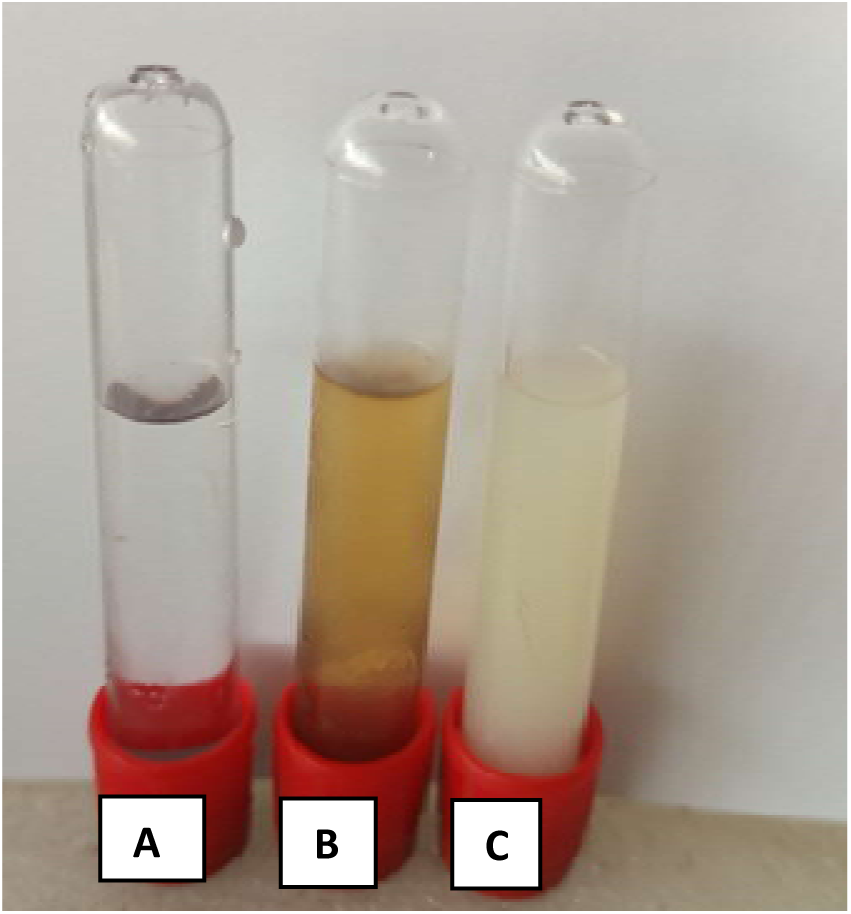
Zinc nitrate hexahydrate solution (A), aqueous extract of *A.citratum* (B) and ZnONPs solution (C)

**Figure 3.**
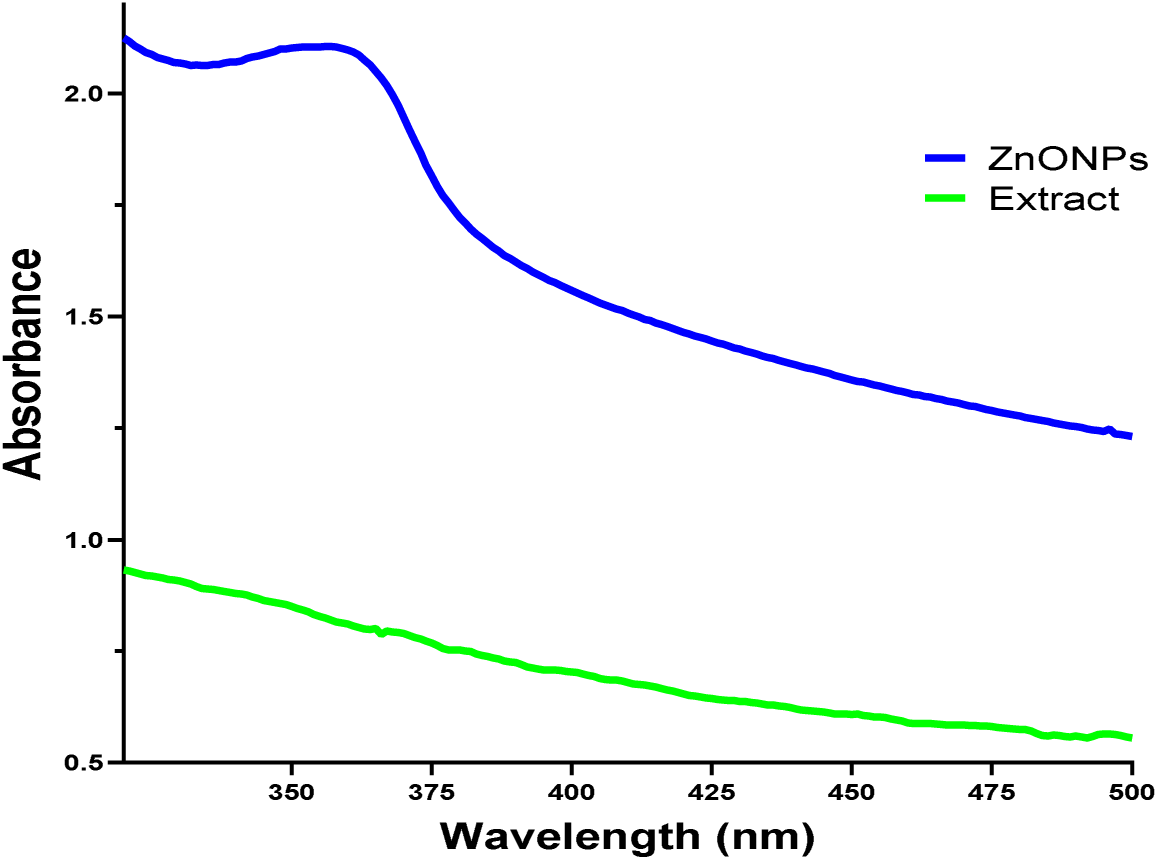
UV/Vis spectrum of ZnONPs with *A.citratum* seed extract

#### UV-Visible spectrophotometry

Figure 15 shows the UV-Vis absorption spectrum of ZnONPs from the aqueous extract of *A. citratum* seeds. It was recorded for samples in the 300-500 nm range. The spectrum showed a plasmon resonance band at 360 nm corresponding to the characteristic band of zinc oxide nanoparticles [16].

### Characterization of zinc oxide nanoparticles

#### Fourier transform infrared spectroscopy (FTIR)

FTIR shows the presence of functional groups from the aqueous extract on the oxide surface. It also suggests that the formation of ZnONPs is due to the interaction of secondary metabolites with zinc nitrate. Figure 4 shows the FTIR spectra of ZnONPs and aqueous extract in the 1000-5000 cm^−1^ range. The spectra show the elongation vibrations of O-H and N-H bonds at 3358 cm, medium-intensity bands at 1028 and 1315 characteristic of C-O and C-N elongation vibrations respectively, and low-intensity bands at 809, 491 and 435cm^−1^characteristic of Zn-O elongation vibrations. Sodium hydroxide was responsible for formation ZnO and the secondary metabolites allowed for the stabilization of the nanoparticles. The functional groups of the metabolites were observed as bands corresponding to various stretching modes. Similar results were observed during the green synthesis of zinc oxide 18using aqueous extracts of *Elaeagnus angustifolia* L. and *Deverra tortuosa* leaves [17, 20].

**Figure 4.**
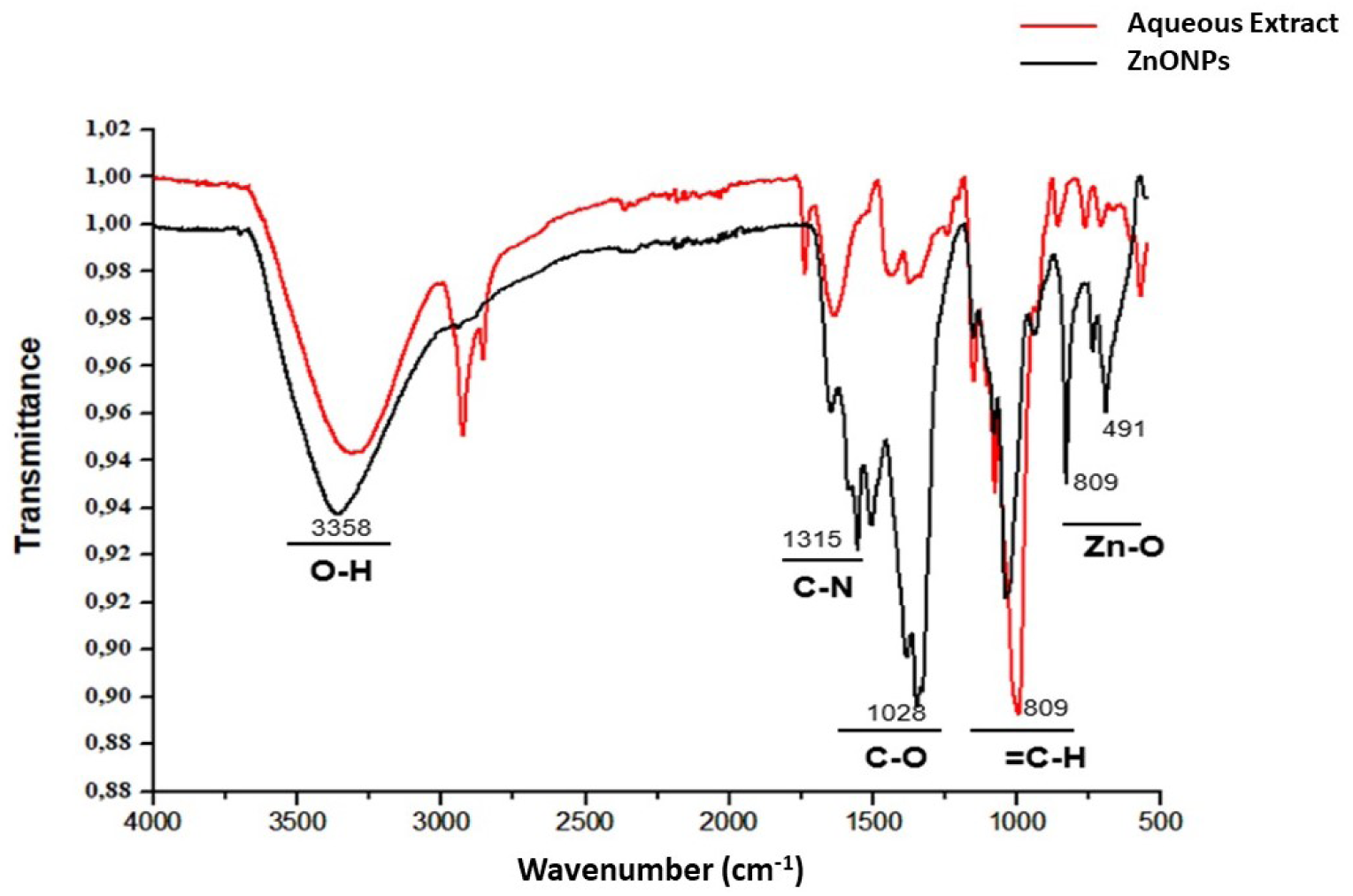
FTIR spectrum of ZnONPs with *A. citratum* seed extract

**Figure 5.**
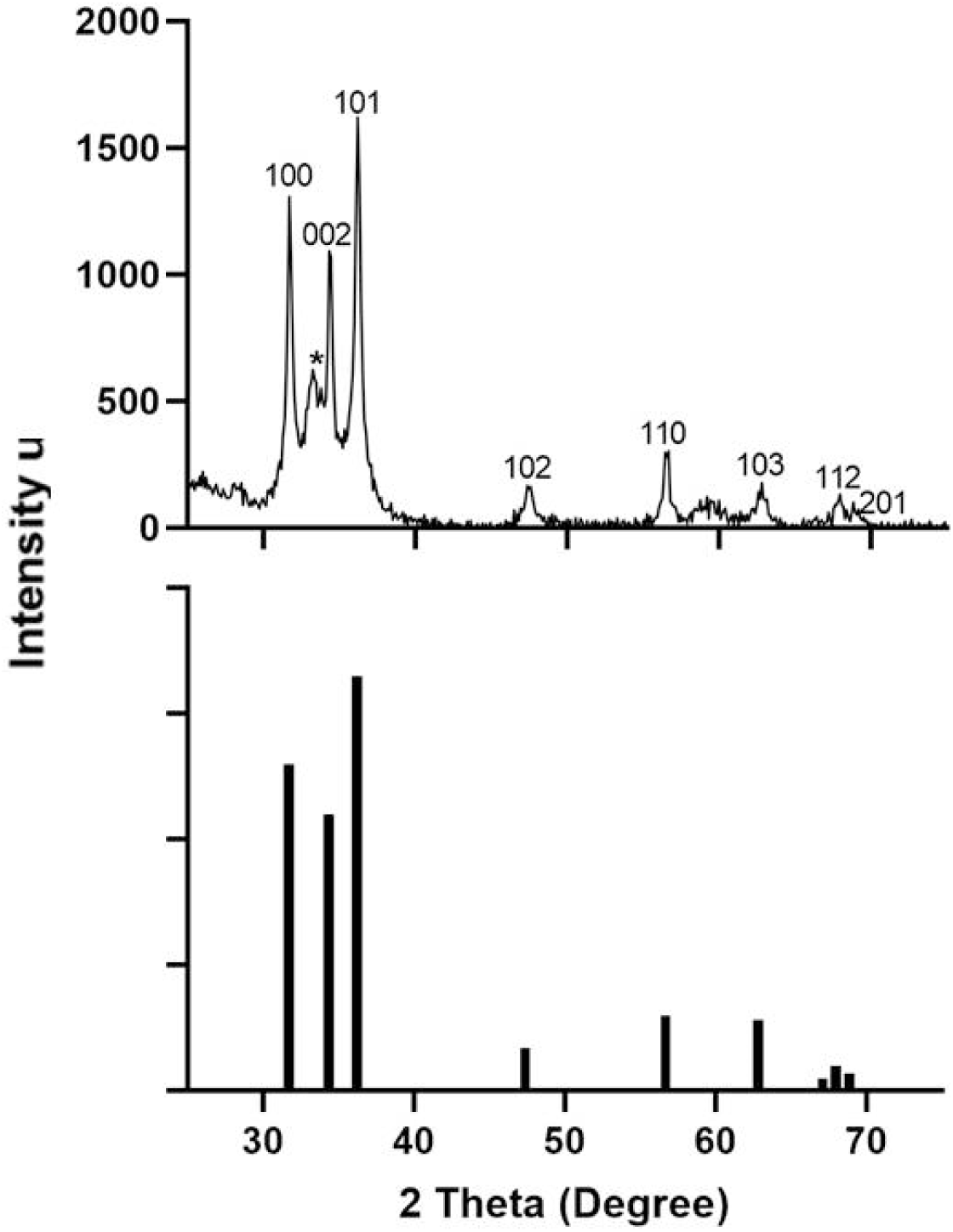
a) PXRD of ZnONPs with *A. citratum* seed extract b) bar diffractogram of wurtzite ZnO (ICDD # 98-002-9272).

#### Powder X-ray diffraction (PXRD)

The diffractogram of ZnONPs indicatedhkl crystalline planes (001), (002), (101), (102), (110), (103), (112) and (201) corresponding to ZnO by comparison with The International Centre for Diffraction Data(ICDD98-002-9272) of ZnO (wurtzite). Undetermined Zn crystallographic phases are observednear 33°.The grain size of 20 nm was determined based Scherrer equation using the hkl plane (101) with the highest intensity.

#### Particles shape, morphology and elemental composition

Synthesized ZnONPs shape and morphology were investigated using scanning electron microscopy (SEM) and are depicted in Figure 6A. The micrograph shows large blocks when isolated from the aqueous dispersion. Transmission electron microscopy (TEM) allowed the visualization of individualZnONPs (Figure 6B) after drops of the dispersion were placed on the sample holder.

**Figure 6.**
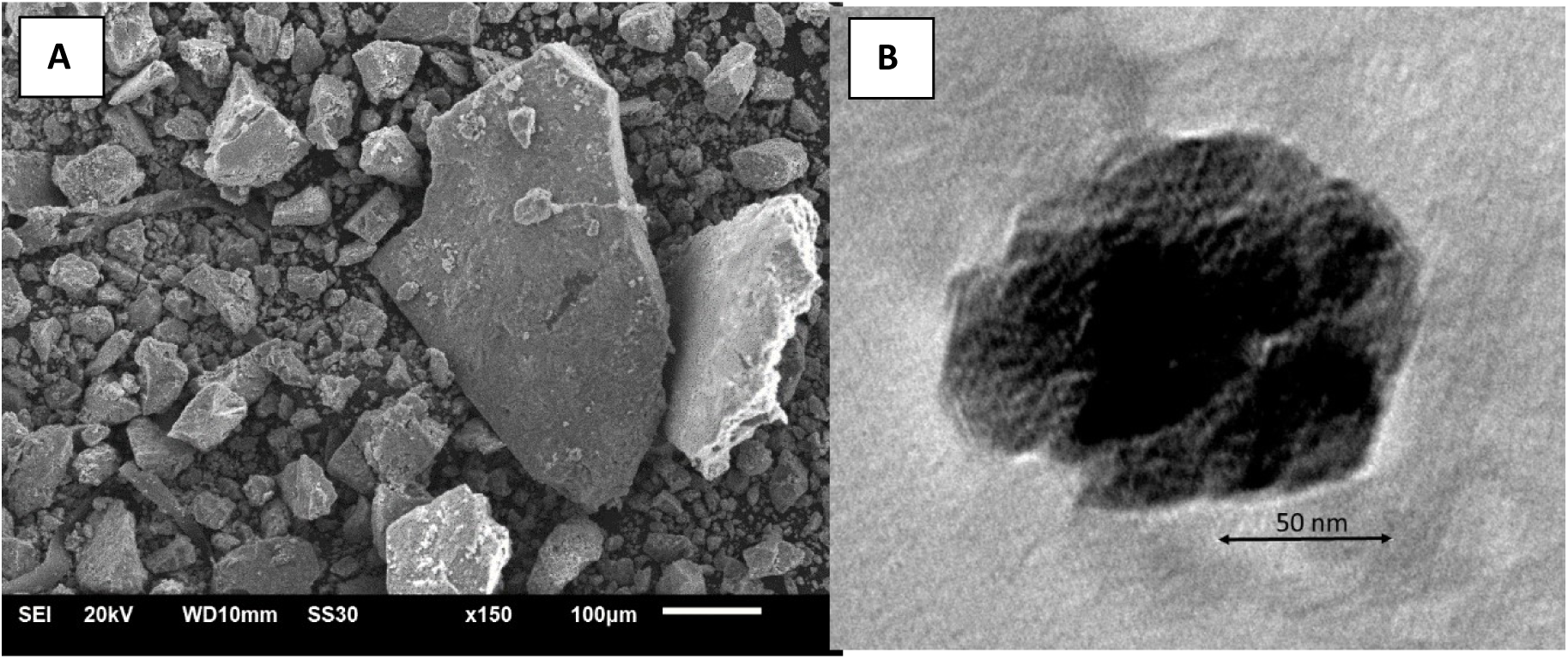
SEM (A) and TEM (B) micrographs of ZnONPs with *A. citratum* seed extract

As for the elemental composition (Table 2), the EDX spectrum depicts three main peaks. Two of those found at 1keV and 8.6 correspond to Zn while the third peak at 0.5 keV corresponds to oxygen (Figure 7, table 2). This result confirms the previous findings by FTIR which has revealed Zn-O vibration band and is in accordance with the results obtained during the synthesis of ZnONPs using *Geranium wallichianum* leaves and orange fruit peel extracts [19].

**Figure 7.**
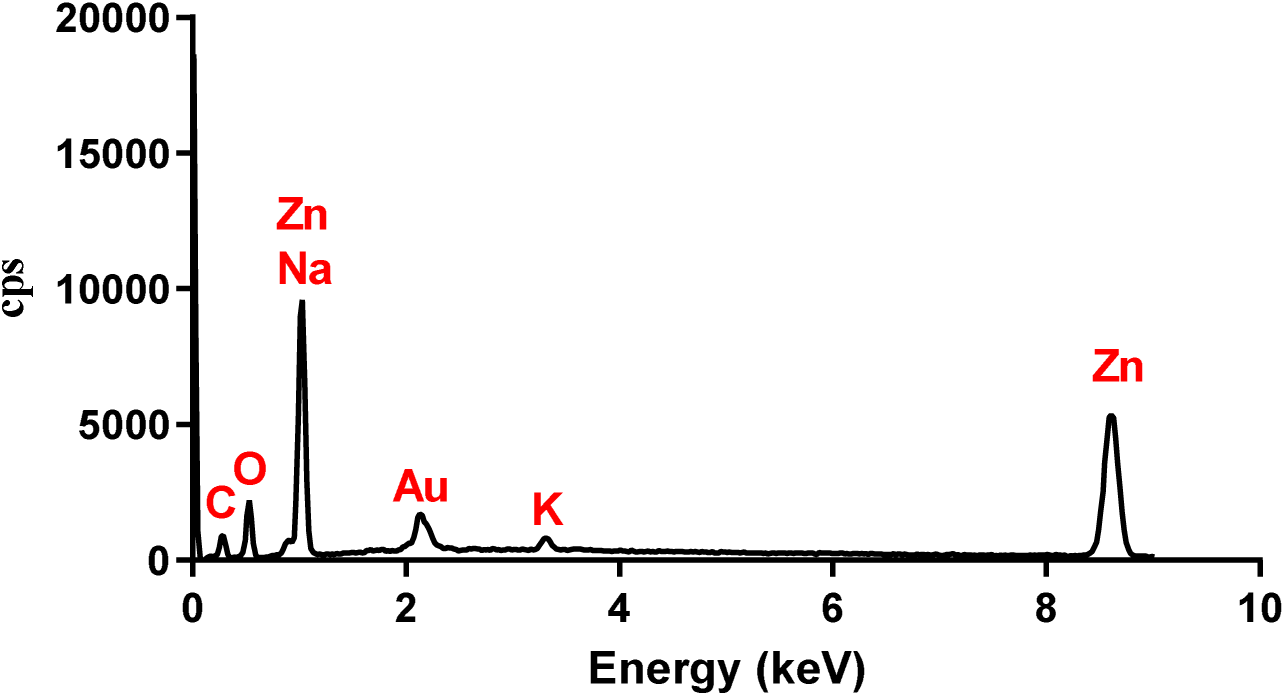
EDx spectrum of ZnONPs main elements

### Acute Toxicity

#### Effects of ZnONPs on clinical parameters

Table 3 shows the clinical parameters follow-up of rats 14 days after administration of a sample of ZnONPs at 2000mg/kg. No signs of toxicity were recorded, and no deaths were noted. The LD_50_ of ZnONPs is probably above 2000 mg/kg, therefore, other toxicological tests must be conducted to determine it.

**Table 3.** Monitoring of rats’ clinical parameters.

| Parameters | Control batch | Test batch (2000 mg/Kg) |
| --- | --- | --- |
| Morbidity | A | A |
| Aggressiveness | A | A |
| Stool appearance | N | N |
| Skin appearance | N | N |
| Eye color | N | N |
| Pain sensitivity | N | N |
| Tremor | A | A |
| Salivation | A | A |

A:absent; N: normal

#### Effect of ZnONPs on rats’body mass

Figure 8 shows the evolution of rat weights as a function of time, showing no significant difference (p > 0.05) in the evolution of treated batch weights compared with the control batch.

**Figure 8.**
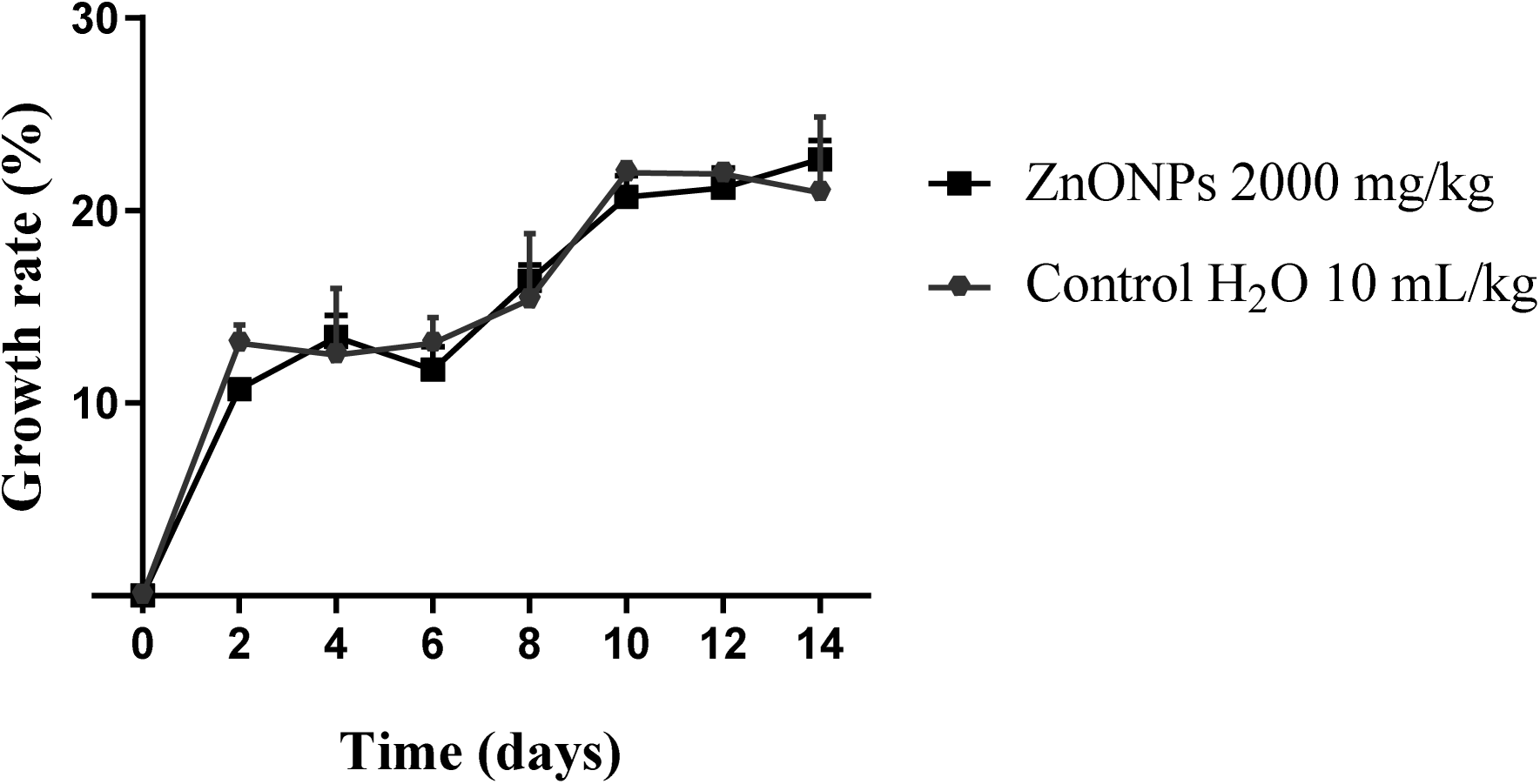
Timely evolution of animals’ weight. ZnONPs: zinc oxide nanoparticles with *A. citratum* seed extract. Results are presented as mean ± SEM (n = 3). No significant difference was observed between the two batches (p >0.05).

#### Effects of ZnONPs on the relative mass of some organs

The vital organs (liver, kidneys, heart, lungs, and spleen) were weighed on day 14, after humanly sacrificing the rats, and their relative mass was calculated (Figure 9). The comparative analysis of the relativeorgan weight showed no significant difference (p > 0.05) between the two batches.

**Figure 9.**
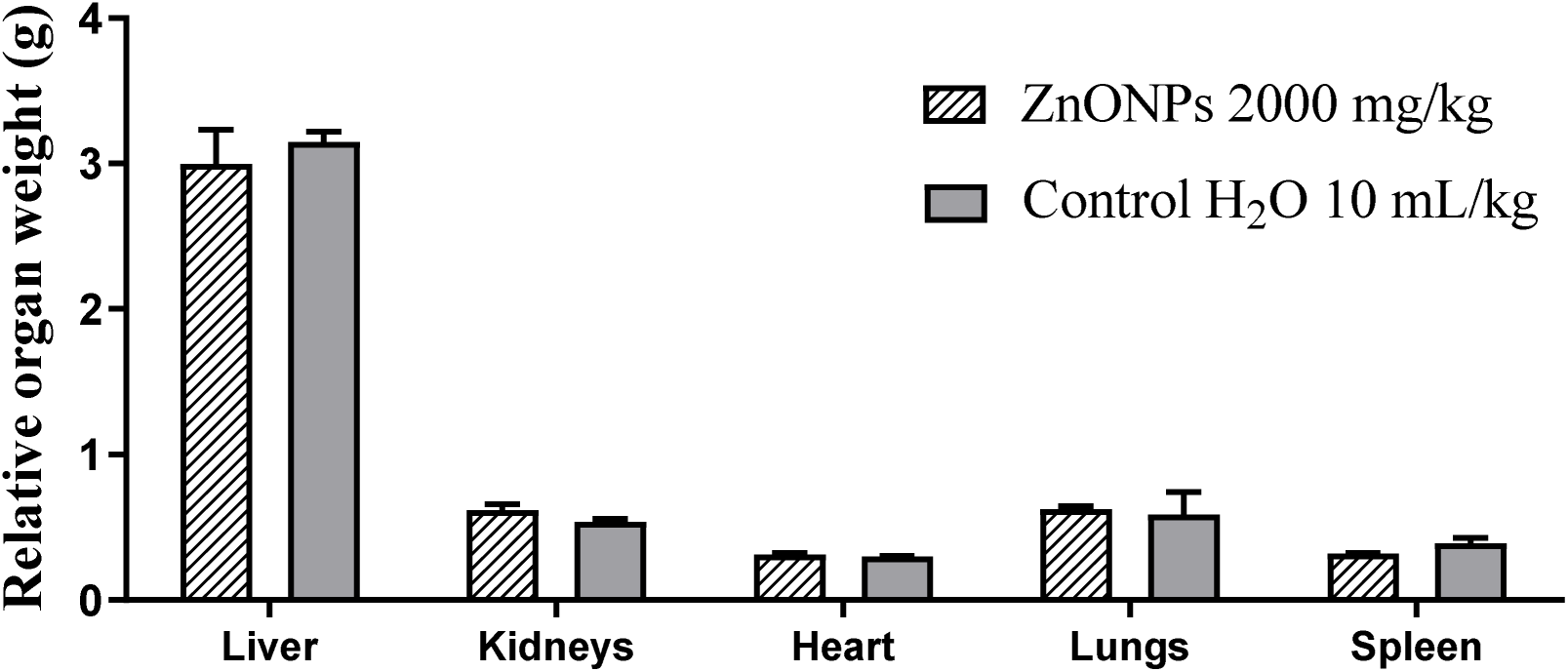
Relative mass of vital organs after 14 days. ZnONPs: zinc oxide nanoparticles with *A. citratum*seed extract. Results are presented as mean ± SEM (n = 3). No significant differences were observed between the groups.

### Effect of the nanoparticlesfrom*A. citratum* on biochemical parameters

#### Effects of ZnONPs on transaminases

Comparative analysis of the two biochemical parameters characteristic of liver function (ALT/AST) showed no significant difference in serum ALT values, while administration of the 2000 mg/kg dose of ZnONPs resulted in a slight increase (p<0.05) in AST compared with the control batch (Figure 10).

**Figure 10.**
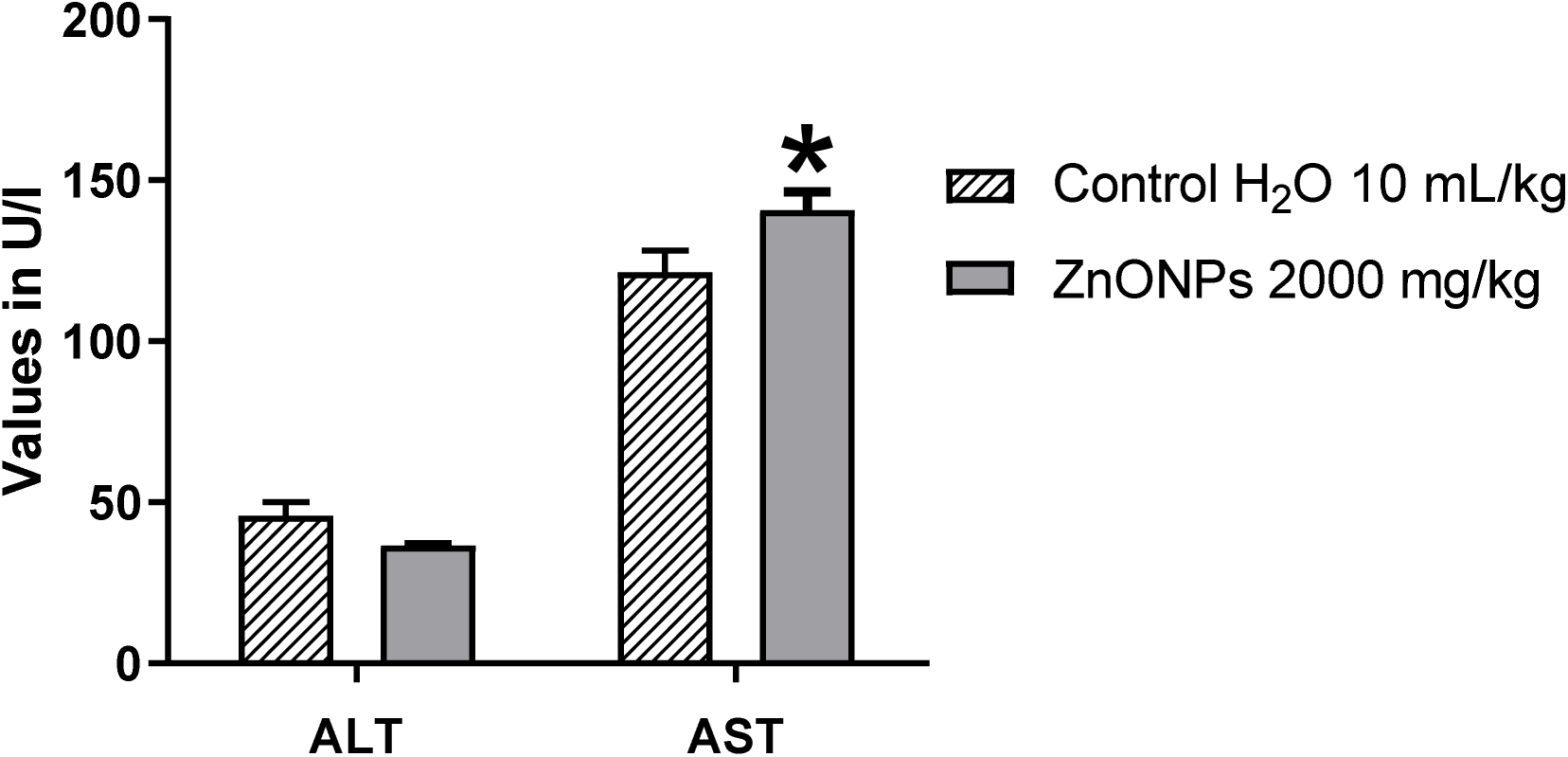
Comparison of serum ALT and AST values of rats after 28 days. Results are presented as mean±SEM (n = 3). (*) significant difference between the two batches (*p* < 0.05).

#### Effects of ZnONPs on markers of renal function

The comparative analysis of two biochemical parameters characteristic of renal function showed no significant difference in urea levels (p = 0.2433) (Figure 11) and creatinine levels (p= 0.3394), between rats treated with ZnONPsand the control group (Figure 12).

**Figure 11.**
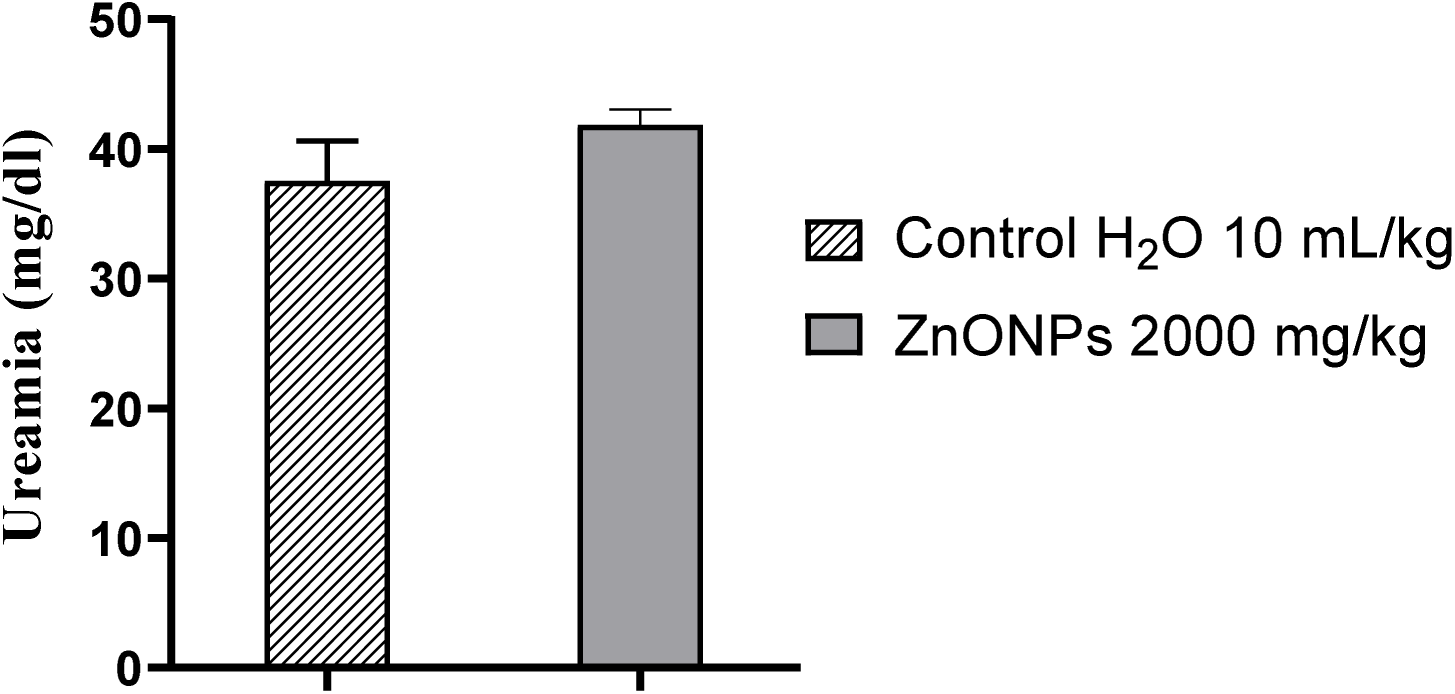
Blood urea levels after 28 days. ZnONPs: zinc oxide nanoparticles with *A. citratum* seed extract. Results are presented as mean ± SEM (n = 3). No significant difference was observed between the two batches (p >0.05).

**Figure 12.**
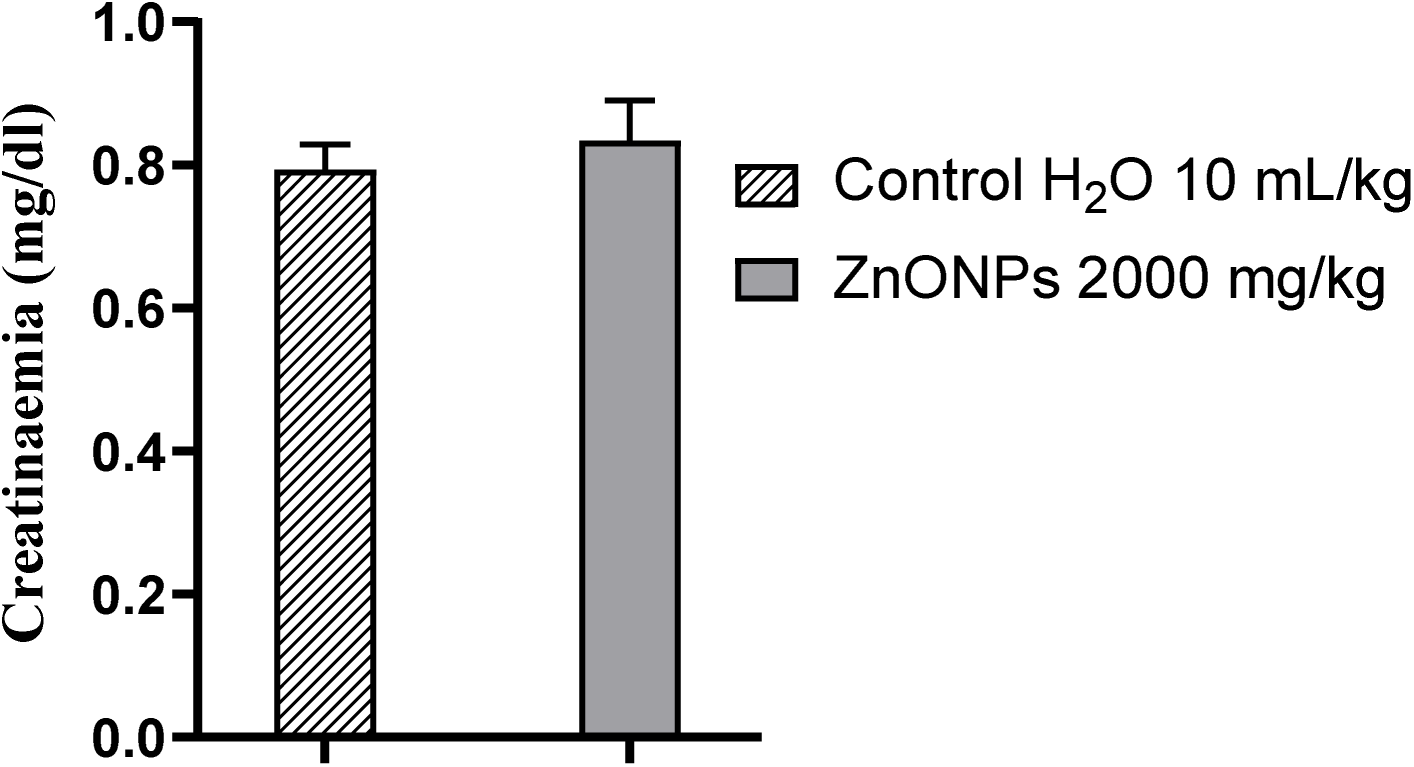
Blood creatinine levels after 28 days. ZnONPs: zinc oxide nanoparticles with *A. citratum* seed extract. Results are presented as mean ± SEM (n=3). No significant difference was observed between the two batches (p > .05).

### *In vitro* anti-inflammatory activity

#### Heat-induced denaturation of egg albumin

A comparative analysis of the information in this table shows that the anti-inflammatory effect of zinc oxide nanoparticles evaluated against egg albumin denaturation showed a maximum inhibition percentage of 87 % at a concentration of 1000 µg/mL; diclofenac used as the standard drug showed an inhibition of 84 % at 600 µg/mL and the aqueous extract was 65% at a concentration of 600 µg/mL (Figure 13).

**Figure 13.**
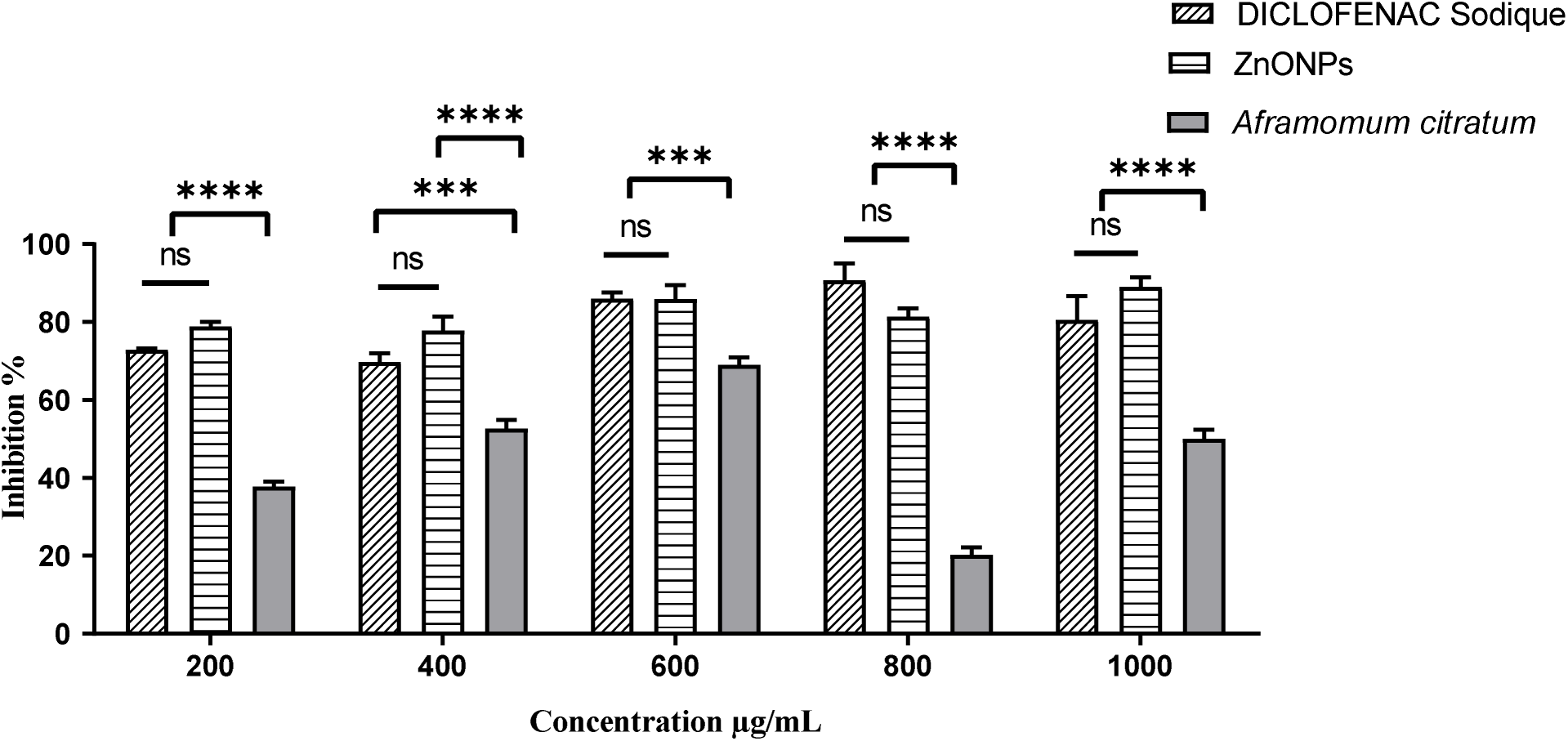
Inhibition rate of albumin denaturation by ZnONPs and *Afromomum citratum* extract. ns = non significative, (***) p < 0.001, (****) p < 0.0001.

#### Effect of ZnONPs on red blood cell membrane stabilization

The test is based on the stabilization of erythrocytes after induction of hemolysis by a hypotonic solution at elevated temperature. The percentage of hemolysis inhibition obtained after incubation with a volume of ZnONPs, aqueous extract and diclofenac sodium (dissolved in NaCl 0.9%) used as reference drug showed that from the results obtained in the hemolysis inhibition histogram (Figure 14), ZnONPs presented a maximum percentage of 77.22% at 600µg/mL, aqueous extract 50% at 400 and 800µg/ml and diclofenac 64% at a concentration of 600µg/mL.

**Figure 14.**
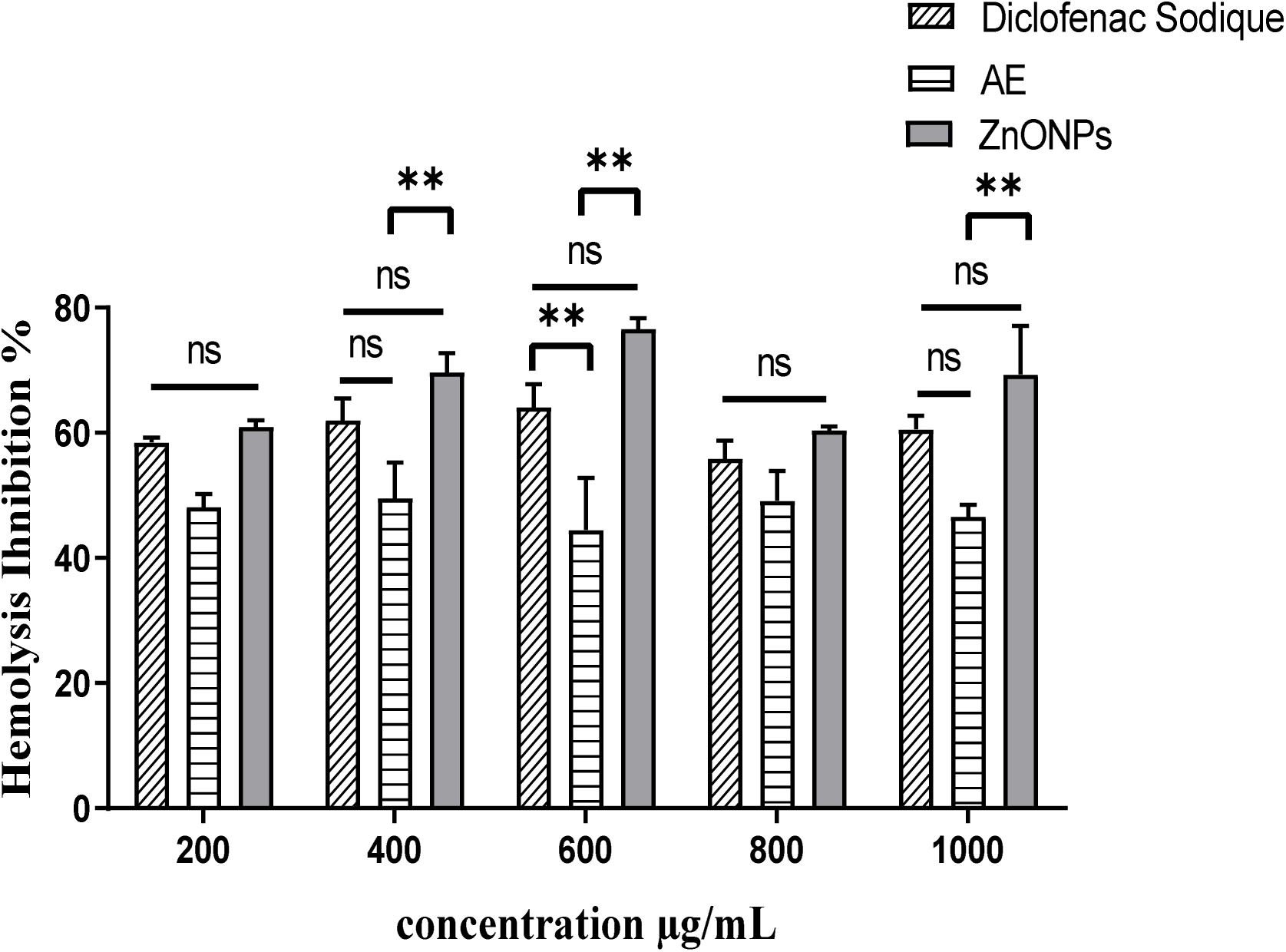
Histogram of percent inhibition of hemolysis by ZnONPs, aqueous extract and diclofenac sodium. ns = non significative, (**) p < 0.01.

#### *In vivo* anti-inflammatory activity on rat plantar edema induced by carrageenan

A comparative analysis of this table reveals that oral administration of ZnONPs resulted in significant inhibition of paw edema compared with the control. The percentages of maximum inhibition obtained were at 30min (53 %), 1st hour (45), 2nd hour (58 %) and 6^th^ hour (79 %) for 0.6, 0.8 and 1mg/kg (body weight) respectively. Diclofenac used as a standard drug had a significant percentage inhibition (%) against carrageenan-induced paw oedema at the 3^rd^ hour (65 %) (Figure 15).

**Figure 15.**
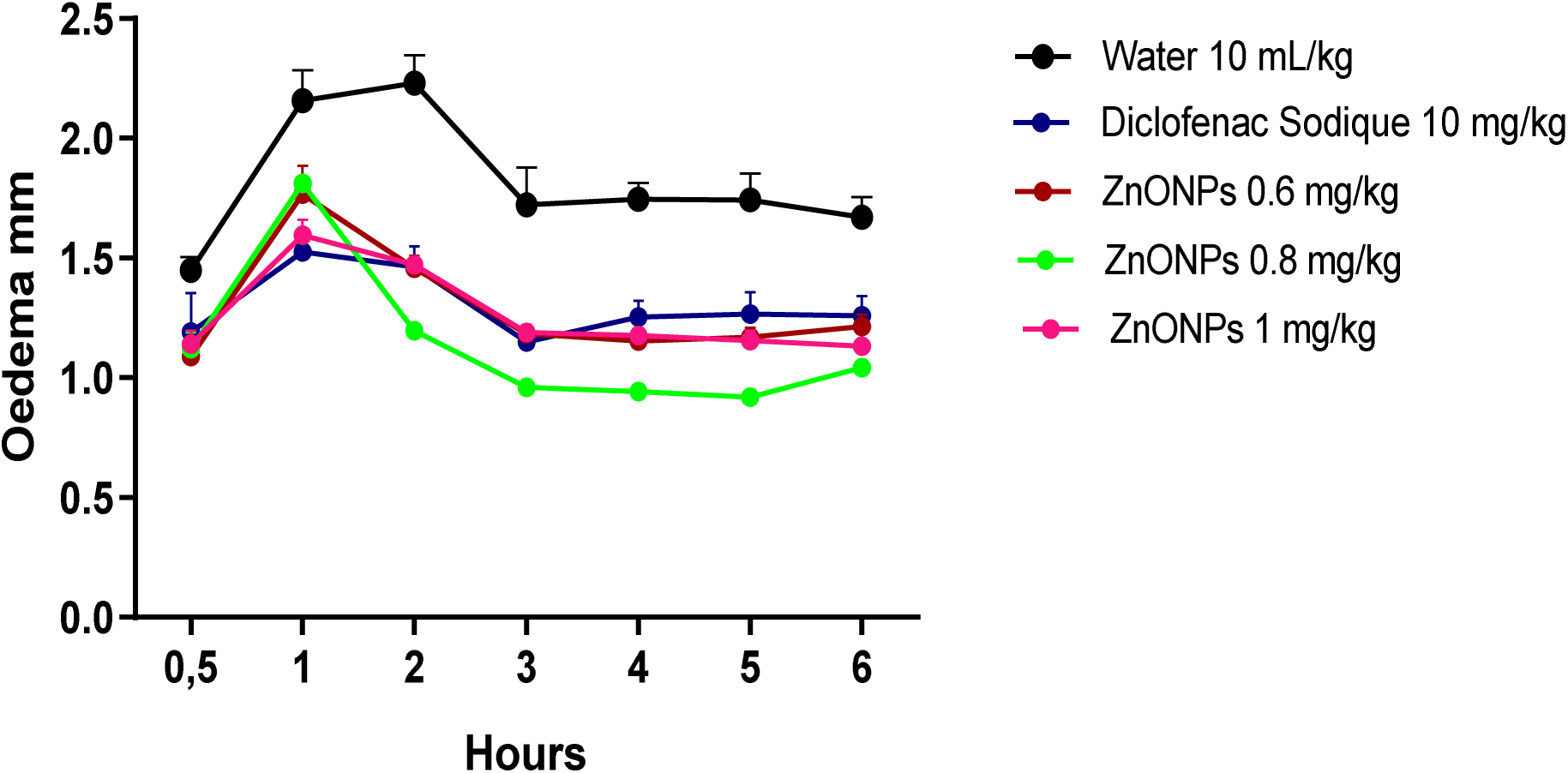
Timely evolution of plantar volume in rats with carrageenan-induced edema, differences are not significative between ZnONPs and diclofenac.

## Discussion

The aqueous extraction of *Aframomum citratum* seeds presented a yield of 9 %, a result close to the value of Nguele *et al.,* [21], who worked on the aqueous extraction of *A. citratum* seeds and obtained a yield of 10 %. The phytochemical composition of the aqueous extract of *A. citratum* seeds revealed the presence of alkaloids, flavonoids, phenols, tannins and anthraquinones and the absence of saponins, triterpenes and anthocyanins. These results are in agreement with those of Fankam*et al.,* [22] but differ from the results of Biqiku *et al.* who showed an absence of tannins, phenols, flavonoids [23] and could be explained by the fact that the composition of *A. citratum* seeds or plant drugs in general varies according to parameters such as: the growing medium of the harvesting season, the state of ripeness of the plant, the drying time, the different extraction methods used. The presence of chemical groups such as phenols, alkaloids, flavonoids, and tannins justify the anti-inflammatory properties and is compatible with the zinc oxide dispersionand their stabilization [24].

Zinc oxide formationis confirmed by the reaction mixture’s color change from brown to pale white and the formation of a turbic, opaque disperson. The secondary metabolites in the zinc oxide solution; the aqueous extract acts as a stabilizing agent. This was confirmed by analysis of the visible spectrum in the 300-500 nm range, which showed a peak at 360 nm that is specific to zinc oxide nanoparticles. These results agree with those of Azim *et al.,* in 2022, who showed an absorbance peak at 360 nm and a pale white color characteristic of zinc oxide nanoparticles [25].

FTIR revealed the compounds present on the nanoparticle surface, as well as the phenolic O-H, C-O, C-N alkane chain and Zn-O functions observed on zinc oxide nanoparticles. These functions agree with the work of Ashwini *et al.*, who considered that phenolic compounds would be involved in the stabilization of nanoparticles [26]. PXRD of *A. citratus* mediated ZnONPs reveals grains with 20 nm size using the 101 hkl plane with the Scherrer equation, smaller than particles obtained by the sol gel method 50 nm and with stirring and ultrasound (80 nm and 100 nm), but similar with methods without agitation (20 and 200 nm) [27, 28].TEM demonstrated the presence of fine particles of a few nanometers in size [29].

The toxicological profile of the ZnONPs solution (administered at a dose of 2000 mg/kg) revealed an absence of probable toxicity. In fact, clinical monitoring, weight evolution, relative comparison of organ weights and the effect of ZnONPs on renal function (urea and creatinine) showed no significant difference between the treated batch and the control batch, while the effect of ZnONPs on liver function (ALT/AST) showed little significant difference between the control batch and the batch treated with ZnONPs, but remained within the range of standards described by Feki *et al.,* in 2021. These results are in agreement with those of Shaban*et al.* [30]. Similarly, no deaths were recorded, suggesting a lethal dose 50 (LD_50_) above the administered dose limit; ZnONPs are therefore not classified by the Organization for Economic Cooperation and Development (OECD) as hazardous products with acute toxicity. These results are in line with those of Natesan *et al*., in 2011, who evaluated the toxicological profile of zinc oxide nanoparticles synthesized from *Elatus anoectochilus* [31] and those obtained by Eyenga *et al.*, who demonstrated that the aqueous extract of *A. citratum* at 2000 mg/kg showed no signs of toxicity [32].

Protein denaturation is one of the causes of complications in inflammatory diseases. As part of the investigation into the mechanism of anti-inflammatory activity *in vitro*, the inhibitory capacity of zinc oxide nanoparticles, aqueous extract of *A. citratum* seeds and diclofenac sodium used as a standard drug were studied against heat-induced egg albumin. Our results showed that at the different concentrations studied, ZnONPs exhibited a high anti-denaturing activity on egg albumin not significativelly different compared diclofenac sodium. The activity of both the nanoparticles and the reference is significativelly higher compared to the plant extract. These results agree with those obtained by Ovia *et al.,* [33].

Concentrations of ZnONPs, aqueous extract and diclofenac sodium, used as standard medication, were tested for their anti-inflammatory efficacy via the red cell membrane stabilization test, as no hemolytic effect was observed. Inflammation is the protective response of living tissue against microbial infections, chemical and physical attack. During the anti-inflammatory process, the release of lysosome constituents such as neutrophils, bactericidal and fungicidal enzymes is suppressed, leading to the release of extracellular components [34]. For these reasons, stabilization of the lysosome membrane is essential to constrain the inflammatory response. In the present work, ZnONPs showed remarkable anti-hemolytic activity compared with aqueous extract and not significativelly different compared to diclofenac sodium; maximum inhibition was recorded at 600 µg/mL. Also, ZnONPs synthesized from *A. citratum* seed extract to inhibit inflammatory responses. They could be a lead actinic compound for the design of an alternative medicine for the management of inflammatory diseases. These results agree with those of Manasa *et al.,* in 2021 [35].

Under experimental conditions, carrageenan induced edema with low inhibition of inflammation during the 1st hour but increased after the 2nd hour. Carrageenan induces local inflammation when injected into the rat’s hind leg due to tissue damage. Its injection triggers the release of several chemical mediators responsible for the inflammatory process. Carrageenan-induced edema is a triphasic response involving the release of various mediators such as histamine and serotonin in the first phase (1st phase); the second phase is mediated by the release of kinin (2nd phase), and the third phase is attributed to prostaglandins and cyclooxygenase products lasting from 3 to 6 hours. Statistical analysis demonstrated significant inhibition by ZnONPs of edema during all three phases. The inhibitions of the nanoparticles are not significatively different compared to the drug at all concentrations. Both responses of the nanoparticles and the diclofenac are significatively higher compared to negative control. Maximum inhibition was observed at a dose of 1 mg/kg (79 %). These results agree with those of Sulaiman *et al*. [36,37]. The authors also suggested that ZnONPs may interfere with the release of acute and chronic inflammatory mediators and that the persistent anti-inflammatory activity may be due to the improved permeability and retention effect of zinc oxide nanoparticles in the edema region reported by Moldovan *et al.* [38].

## Conclusion

The work carried out involved the synthesis, characterization and evaluation of the anti-inflammatory activities of ZnONPs obtained from *Aframomum citratum* seeds, as well as the assessment of their acute and hemolytic toxicity. The synthesized ZnONPs were confirmed after visual observation by ultraviolet visible spectrophotometry, the infrared performed showed the presence of functional groups on their surface, their elemental mapping as well as their shape and size were carried out. They showed no signs of harmful toxicity in relation to the toxicity tests carried out. Furthermore, ZnONPs obtained from the extract proved more effective than the aqueous extract in inhibiting heat-induced egg albumin, carrageenan-induced edema, and red blood cell membrane stabilization.

At the end of this study, the observations and results allow us to conclude that ZnONPs obtained from *A. citratum* seeds possess a more potent anti-inflammatory activity than the aqueous extract and are a probable alternative in the treatment of inflammatory pathologies.

## Ethics Statement

The animals were examined and adapted to the new environmental conditions for a week before the formal experiment. All experimental procedures were in strict compliance with theapproved protocol by the Institutional Ethic Committee For Human Research of the University of Douala (Protocol approvalnumber 3544 CEI-Udo/03/2023T).

## Acknowledgements

FEM thank the DAAD for a generous Visiting Professor Fellowship (grant no. 57588364).

## Data availability

The authors declare that the data supporting the findings of this study including raw data files are available from the corresponding author upon reasonable request.

## Statement on animal rights

On behalf of all authors, the corresponding author affirms that animal rights were upheld in the study

## Declarations

All authors approve the submission

## Author Contribution

Francois Eya’ane Meva, Christoph Janiak; Writing – review & editing, Conceptualization, Supervision, Investigation, Funding acquisition. Denise Murielle Nga Essama, Edvige Laure Nguemfo, Hans-Denis Bamal, Agnes Antoinette Ntoumba, Phillipe Belle Ebanda Kedi, Thi Hai Yen Beglau. Writing – review & editing, Conceptualization, Visualization, Intrumentation. Alex Kevin Tako Djimefo, Annie Guilaine Djuidje, Geordamie Chimi Tchatchouang, Chick Christian Nanga, Gildas Fonye Nyuyfoni, Armel Florian Tchangou Njimou, Danielle Ines Madeleine Evouna, Armel Ulrich Mintang Fongang. Writing – review & editing, Investigation, Data curation, Animal models, Instrumental support.

